# pTSara-NatB, an improved N-terminal acetylation system for recombinant protein expression in *E. coli*

**DOI:** 10.1101/331355

**Authors:** Matteo Rovere, Alex E. Powers, Dushyant S. Patel, Tim Bartels

**Affiliations:** Ann Romney Center for Neurologic Diseases, Brigham and Women’s Hospital and Harvard Medical School, 60 Fenwood Road, 02115-6128 Boston, MA, United States

**Author notes:** (TB).

## Abstract

N-terminal acetylation is one of the most common post-translational modifications of the eukaryotic proteome and regulates numerous aspects of cellular physiology, such as protein folding, localization and turnover. In particular α-synuclein, whose dyshomeostasis has been tied to the pathogenesis of several neurodegenerative disorders, is completely N^α^-acetylated in nervous tissue. In this work, building on previous reports, we develop and characterize a bacterial N-terminal acetylation system based on the expression of the yeast N-terminal acetyltransferase B (NatB) complex under the control of the P_BAD_ (L-arabinose-inducible) promoter. We show its functionality and the ability to completely N^α^-acetylate our model substrate α-synuclein both upon induction of the construct with L-arabinose and also by only relying on the constitutive expression of the NatB genes.

## Introduction

Protein N^α^-acetylation, or N-terminal acetylation, is one of the most common post-translational modifications (PTMs) of the eukaryotic proteome, with a vast majority of all N-termini (∼80%) bearing this moiety. The reaction is catalyzed by a class of enzymes, N-terminal acetyltransferases (NATs), of which six (NatA to NatF) have to-date been discovered in humans and one (NatG) has been identified in *Arabidopsis thaliana*, with no human ortholog [1]. These enzymes mediate the transfer of an acetyl group from acetyl-CoA to the positively charged N-terminus of the protein. Their activity often requires the formation of a complex with the ribosome, mediated by one or two auxiliary, ribosome-anchoring, subunits, which provide scaffolding for the catalytic subunit and, in some cases, also regulate its substrate specificity [2,3]. N^α^-acetylation thus occurs usually [1,4] in a co-translational fashion, with the acetyl moiety being added to the nascent polypeptide chain [5,6]. Different enzymes of the NAT family will show different specificities for the polypeptidic substrates to be N-terminally acetylated, based on the first 2-4 amino acids of the nascent chain [1]. The role of N-terminal acetylation varies wildly from protein to protein and organism to organism, but it has been shown to be central to protein homeostasis and cellular physiology, regulating protein half-lives, protein-protein interactions, subcellular localization, folding and aggregation [1]. α-synuclein (αSyn) is one of proteins for which the effects of N-terminal acetylation have been shown to be central to its physiology and pathology. αSyn is a small protein (140 aa, 14.6 kDa) ubiquitously and abundantly expressed in nervous tissue [7,8]. While its exact function is still unclear, it has long been associated with the regulation of synaptic activity and neurotransmitter release [7,9]. Most importantly, both genetic and histopathologic evidence have tied it to the pathogenesis of a class of diseases known as synucleinopathies [10], including Parkinson’s Disease, the second most common progressive neurodegenerative disorder [11]. The totality of αSyn in human tissue has been shown to be N^α^-acetylated [12,13] and a number of studies have highlighted the role of this PTM in the modulation of αSyn’s lipid binding, aggregation, oligomerization and helical propensity [14–17]. This is especially important given the ongoing discussion on the structure of native αSyn in a cellular environment, which requires structural studies to be performed on a species as close as possible to the one present in nervous tissue [18].

While the expression of recombinant, N^α^-acetylated proteins is possible in eukaryotic hosts as yeast and insect cells, a prokaryote-compatible N^α^-acetylation system in bacteria (e.g. *E. coli*) would provide a cheaper and easier-to-use alternative. Although NATs are present in bacteria and archaea the occurrence of N-terminal acetylation is much lower and the NATs’ specificity and regulation are not well-characterized [19]. One approach has been to co-express yeast NATs along with the target protein in bacteria and it has been applied successfully to most of the NATs’ substrates [20,21]. While promising, this method has some shortcomings. The overexpression of both NATs and the target protein under the same inducible promoter does not ensure the proper folding and assembly of the NAT complex before the expression of the N^α^-acetylation target begins, which, given the co-translational nature of the PTM, can lead to N^α^-acetylated/non-N^α^-acetylated mixtures [21,22]. We have thus developed an improved N^α^-acetylation system, pTSara-NatB, under the control of a P_BAD_ promoter [23] and tested its performance using αSyn as a model substrate.

## Materials and Methods

### Materials

All materials were obtained from Sigma-Aldrich (St. Louis, MO), unless otherwise noted.

### Molecular Cloning

pTSara was a gift from Matthew Bennett (Addgene plasmid # 60720) [24]. pNatB (pACYCduet-naa20-naa25) was a gift from Dan Mulvihill (Addgene plasmid # 53613) [20]. Mutagenesis of the pNatB construct to correct the A2520G mutation was performed using the QuikChange II site-directed mutagenesis kit (Agilent Technologies, Santa Clara, CA) and primers G2520A FWD 5’-CGTCGTTTGAATGTATGAATCGATCATTCCTTCACCAAC-3’ and G2520A REV 5’-GTTGGTGAAGGAATGATCGATTCATACATTCAAACGACG-3’; to insert a PvuI restriction site in pTSara, upstream of the T7Te terminator, the primers PvuI FWD 5’-TGTGATCCAAGCCAGCTCGATCGCCGTCGGCTTG-3’ and PvuI REV 5’-CAAGCCGACGGCGATCGAGCTGGCTTGGATCACA-3’ were used. The Naa20 insert in pNatB_G2520A was PCR-amplified using the primers Naa20 FWD 5’-TTGGGCTAGCACTAGTTATAAGAAGGAGATATACATATG-3’ and Naa20 REV 5’-ATGCCTGCAGGTCGACCTAAAATGAAACATCAGCTGG-3’ and inserted into pTSara_PvuI (linearized with SpeI/SalI) using the In-Fusion HD Cloning Kit (Takara Bio, Mountain View, CA). The Naa25 insert in pNatB_G2520A was PCR-amplified using the primers Naa25 FWD 5’-TTTTTTGGGCTAGCGAGCTCTATAAGAAGGAGATATACATATGCGTCGTTCTGGGAGTAAAGAATC-3’ and Naa25 REV 5’-ATCCAAGCCAGCTCGATCGCTAAAATTTTACAAATTTTGGAAGCTTGCT-3’ and inserted into pTSara_PvuI-Naa20 (linearized with SacI/PvuI) using the In-Fusion HD Cloning Kit (Takara Bio, Mountain View, CA). Cloning of pTSara_PvuI-Naa20-Naa25 (pTSara-NatB) and of all of the cloning intermediates was confirmed by DNA sequencing (Molecular Biology Core Facilities, Dana-Farber Cancer Institute) and restriction analysis.

### αSyn Expression and Purification

pET21a-alpha-synuclein was a gift from the Michael J. Fox Foundation MJFF (Addgene plasmid # 51486). BL21(DE3) *E. coli* (New England Biolabs, Ipswich, MA) were freshly co-transformed with pET21a-alpha-synuclein and pTSara-NatB and selected on ampicillin-(amp) and chloramphenicol-(cam) supplemented LB-agar plates. Cultures were grown in LB+amp+cam and induced at an OD_600_ of 0.5-0.6 with 0.2% (m/v) L-arabinose and, after 30 min., with 1 mM isopropyl-β-D-thiogalactopyranoside (IPTG, or with IPTG alone at an OD_600_ of 0.5-0.6). Growth was continued for 4 hrs. at 37°C under shaking. The cell pellet, after being harvested and kept frozen at −20°C overnight, was resuspended in 20 mM Tris buffer, 25 mM NaCl, pH 8.00, and lysed by boiling for 15 min. The supernatant of a 20-min., 20,000xg spin of the lysate was then further processed. The sample was loaded on two 5-mL (tandem) HiTrap Q HP anion exchange columns (GE Healthcare, Pittsburgh PA), equilibrated with 20 mM Tris buffer, 25 mM NaCl, pH 8.00. αSyn was eluted from the columns with a 25-1000 mM NaCl gradient in 20 mM Tris buffer, 1 M NaCl, pH 8.00. For hydrophobic interaction chromatography αSyn peak fractions were pooled and injected on two 5-mL (tandem) HiTrap Phenyl HP hydrophobic interaction columns (GE Healthcare, Pittsburgh, PA), equilibrated with 50 mM phosphate buffer, 1 M (NH_4_)_2_SO_4_, pH 7.40. αSyn was eluted from the columns with a 1000-0 mM (NH_4_)_2_SO_4_ gradient in Milli-Q water. αSyn peak fractions were then pooled and further purified via size-exclusion chromatography on a HiPrep Sephacryl S-200 HR 26/60 column (GE Healthcare, Pittsburgh, PA) using 50 mM NH_4_Ac, pH 7.40 as running buffer. αSyn peak fractions were pooled, aliquoted, lyophilized and stored at −20°C.

### Antibodies

2F12 mouse mAb against human αSyn and Anti-NAT5 mouse mAb against human Naa20 (clone 2C6) were obtained from Sigma-Aldrich (St. Louis, MO) and used, respectively, at 1:10,000 and 1:1000 dilution. Anti-C12orf30 rabbit pAb against human Naa25 was obtained from Abgent (San Diego, CA) and used at a 1:1000 dilution.

### SDS-PAGE and Immunoblotting

Electrophoresis and blotting reagents were obtained from Thermo Fisher Scientific (Waltham, MA), unless otherwise noted. Samples were prepared for electrophoresis by the addition of 4x NuPAGE LDS sample buffer supplemented with 2.5% β-mercaptoethanol and denatured at 85°C for 10 min. Samples were electrophoresed on NuPAGE Novex 4-12% Bis-Tris gels with NuPAGE MES-SDS running buffer and using the SeeBlue Plus2 MW marker. Gels were Coomassie Brilliant Blue- (CBB) stained using GelCode Blue Safe Protein Stain, according to the manufacturers’ protocol, and imaged using a LI-COR Odyssey Classic scanner (LI-COR Biosciences, Lincoln, NE). After the electrophoresis, for immunoblotting, gels were electroblotted onto Immobilon-PSQ 0.2 μm PVDF membrane (Millipore, Billerica, MA) for 1 hr. at 400 mA constant current at 4°C in 25 mM Tris, 192 mM glycine, 20% (v/v) methanol transfer buffer. After transfer, the membranes of gels run with lysate samples were incubated in 4% (m/v) paraformaldehyde in phosphate buffered saline (PBS) for 30 min. at RT, rinsed (3x) 5 min. with PBS and blocked with a 5% milk solution (PBS containing 0.1% (v/v) Tween 20 (PBS-T) and 5% (m/v) powdered milk) for either 1 hr. at RT or overnight at 4°C. After blocking, membranes were incubated in primary antibody in 5% milk solution for either 1 hr. at RT or overnight at 4°C. Membranes were washed (3x) 5 min. in PBS-T at RT and incubated (30 min. at RT) in horseradish peroxidase-conjugated secondary antibody (GE Healthcare, Pittsburgh, PA) diluted 1:10,000 in 5% milk solution. Membranes were then washed (3x) 5 min. in PBS-T and developed with SuperSignal West Dura according to manufacturers’ instructions.

### Mass Spectrometry

Samples were analyzed on an ABI 4800 TOF/TOF Matrix-Assisted Laser Desorption Ionization (MALDI) mass spectrometer (Applied Biosystems, Foster City, CA). Samples undergoing trypsin digestion were incubated overnight in 50 mM NH_4_HCO_3_, 5 mM CaCl_2_, and 12.5 ng·*μ*L^-1^of trypsin, then desalted and concentrated using Millipore C18 ZipTips before spotting. Both trypsin-digested samples and samples for intact mass analysis were prepared for spotting by mixing 0.5 µL of sample with 0.5 µL of α-cyano-4-hydroxy-*trans*-cinnamic acid (10 mg·ml^-1^ in 70% acetonitrile, 0.1% TFA). After drying, samples were rinsed with 0.1% TFA. In addition to external calibration, when measuring intact masses insulin was added as an internal standard, for higher accuracy.

### Growth Curves

Colonies of either singly transformed (pET21a-alpha-synuclein) or co-transformed (pET21a-alpha-synuclein+pTSara-NatB) BL21(DE3) *E. coli* (New England Biolabs, Ipswich, MA) were picked from fresh (<2 weeks) agar-LB+amp or agar-LB+amp+cam plates and inoculated in LB+amp or LB+amp+cam. After 8-10 hrs. of growth, at 37°C under shaking, the cultures, then in their stationary phase, were diluted 1:30 in fresh medium+antibiotic (and 0.2% L-arabinose in one case), aliquoted in 96-well clear sterile plastic plates, sealed with gas-permeable sealing membranes and grown at 37°C under shaking overnight. Absorbance (optical density) at 600 nm (OD_600_) was measured every 15 min. with a Synergy H1 microplate reader (BioTek, Winooski, VT). Data were analyzed with GraphPad Prism 7 (GraphPad Software, La Jolla, CA).

## Results

In uncoupling the induction of the NatB complex and αSyn (or any of NatB’s substrates) two courses of action are possible: either changing the operon regulating the transcription of the NatB genes or the one acting on the SNCA (αSyn) gene. While the authors of the original NatB work suggest [22] and recently implemented [21] an N-terminal acetylation system where the target protein is under a rhamnose-inducible promoter, we decided to redesign pNatB into an arabinose-inducible system. This approach provides two clear advantages. First, using a promoter weaker than the T7/lac of the pET system will dramatically decrease the protein yield (one of the reasons for employing a bacterial expression system in the first place). In addition, the function of the N-terminal acetylation complex can be performed by catalytic amounts of enzyme and, as such, low expression levels should be more than sufficient for the complete modification of the target and, at the same time, pose less of a metabolic burden to the cells. Following the original approach used for pNatB and starting from the bicistronic construct pTSara [24], we cloned both the catalytic, Naa20, and regulatory, Naa25, subunit into pTSara, maintaining the ribosome-binding region of pACYC-Duet-1 (a previously reported missense A-to-G mutation in the Naa25 gene [25] was also corrected), (Fig 1A) and called the construct pTSara-NatB. We then verified the success of the expression by CBB-stained SDS-PAGE and immunoblotting of Naa20 and Naa25 (Fig 1B and C). In addition, we tested the compatibility of pTSara-NatB with the SNCA expression vector (pET21a-alpha-synuclein) by co-transforming and inducing doubly-selected cells containing both plasmids. 0.2% of L-arabinose, which has been shown to promote a robust expression of P_BAD_-regulated genes [23], was used for the induction of the *ara* operon. L-arabinose was added upon reach of a culture density (OD_600_) of about 0.5, 30 min. before the addition of IPTG for pET induction. Both the expression of the NatB subunits and that of the target protein appear to be unaffected by the co-expression and there is no evidence of cross-talk (e.g. αSyn expression upon arabinose addition) between the operons (Fig 1D).

**Fig 1.**
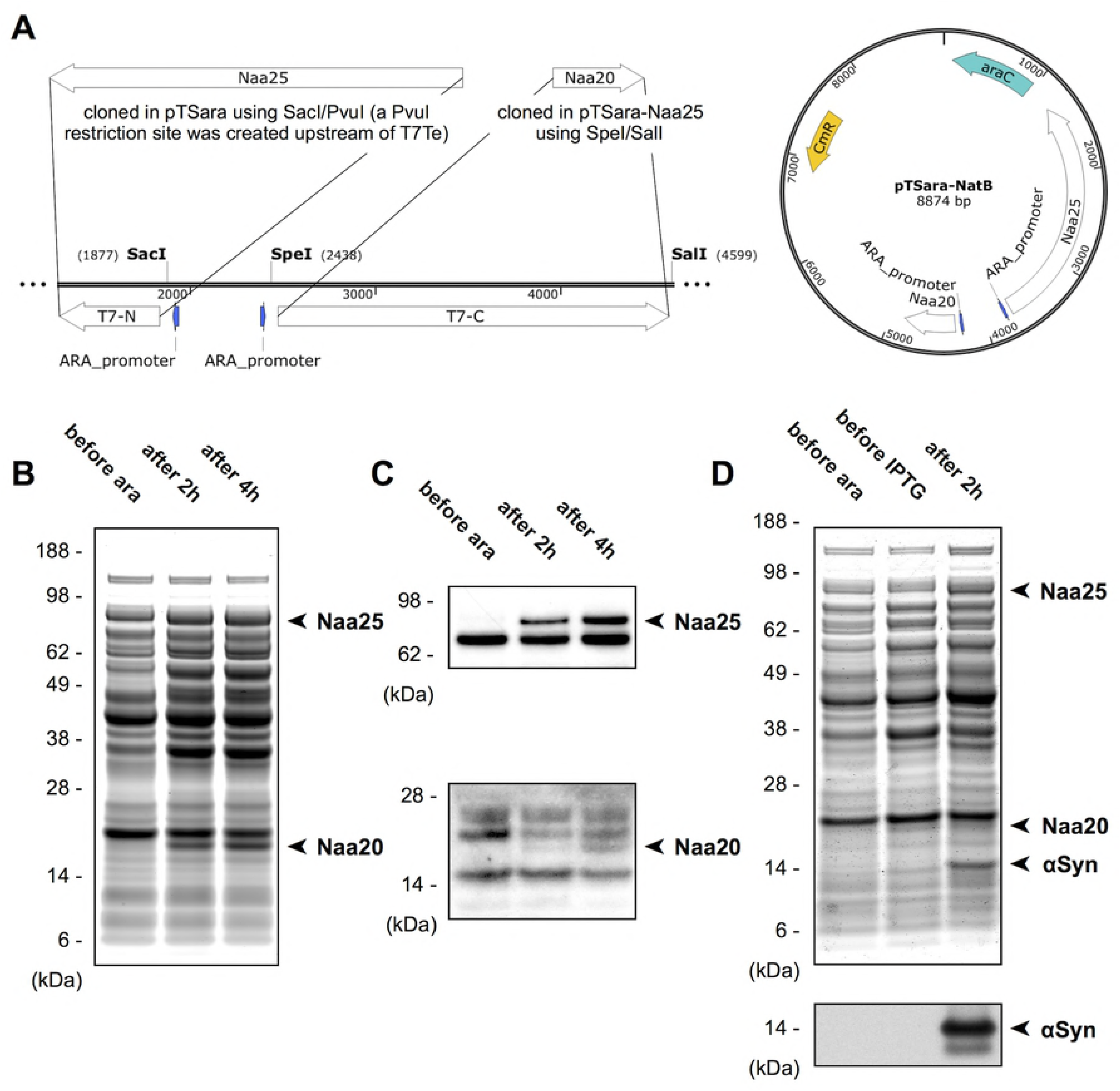
Molecular cloning and characterization of pTSara-NatB. (A) Cloning strategy and plasmid map of pTSara-NatB (B) CBB-stained SDS-PAGE of pTSara-NatB-transformed *E. coli* (PBS-soluble) lysates before and after (2, 4 hrs.) induction with 0.2% L-arabinose. Bands corresponding to the regulatory (Naa25) and catalytic (Naa20) subunits of the NatB complex are marked. (C) Western blots of pTSara-NatB-transformed *E. coli* lysates before and after (2, 4 hrs.) induction with 0.2% L-arabinose. Antibodies to the human homologs of the yeast NatB components were used for detection (*top* Naa25, Anti-C12orf30 1:1000; *bottom* Naa20, Anti-NAT5 1:1000). Non-marked bands are cross-reactive *E. coli* proteins. (D, *top*) CBB-stained SDS-PAGE of pET-alpha-synuclein+pTSara-NatB co-transformed *E. coli* lysates before 0.2% L-arabinose induction (before ara) or 1 mM IPTG induction (before IPTG, added 30 min. after L-arabinose) and 2 hrs. after IPTG induction. Both subunits of the NatB complex and αSyn are marked. (D, *bottom*) αSyn Western blot of co-transformed *E. coli* lysates, in order to confirm the absence of any cross-reactivity between L-arabinose and IPTG induction, 2F12 (1:10,000) was used for αSyn detection.

The N-terminal acetylation efficiency of pTSara-NatB was then tested, using αSyn as a substrate, with the same protocol described before for double transformation and sequential induction. Matrix-Assisted Laser Desorption Ionization-Time Of Flight (MALDI-TOF) Mass Spectrometry (MS) of the purified protein from BL21(DE3) *E. coli* co-transformed with pTSara-NatB and pET21a-alpha-synuclein, either induced or non-induced with L-arabinose, shows, somewhat surprisingly, complete substrate N^α^-acetylation in both cases (Fig 2). However, these results can be easily explained by the fact that complete silencing of the *ara* operon is not attainable by simple absence of the inducer. The catalytic nature of the NatB complex ensures that even small amounts, constitutively expressed, can acetylate efficiently the totality of the target protein. Confirming this mechanistic explanation, addition of D-glucose (0.2%) to the bacterial cultures, which has been shown to reduce the level of non-induced expression of P_BAD_-regulated genes through catabolite repression [23,26], reduced the fraction of N^α^-acetylated αSyn to about 50% (S1 Fig). We also found that such mixtures of N^α^-acetylated and non-N^α^-acetylated αSyn can be resolved by hydrophobic interaction chromatography (S2 Fig). pTSara-NatB thus works as a low-level constitutive expression vector and can potentially be L-arabinose-regulated in the case of difficult substrates (see, *e.g.*, [20]).

**Fig 2.**
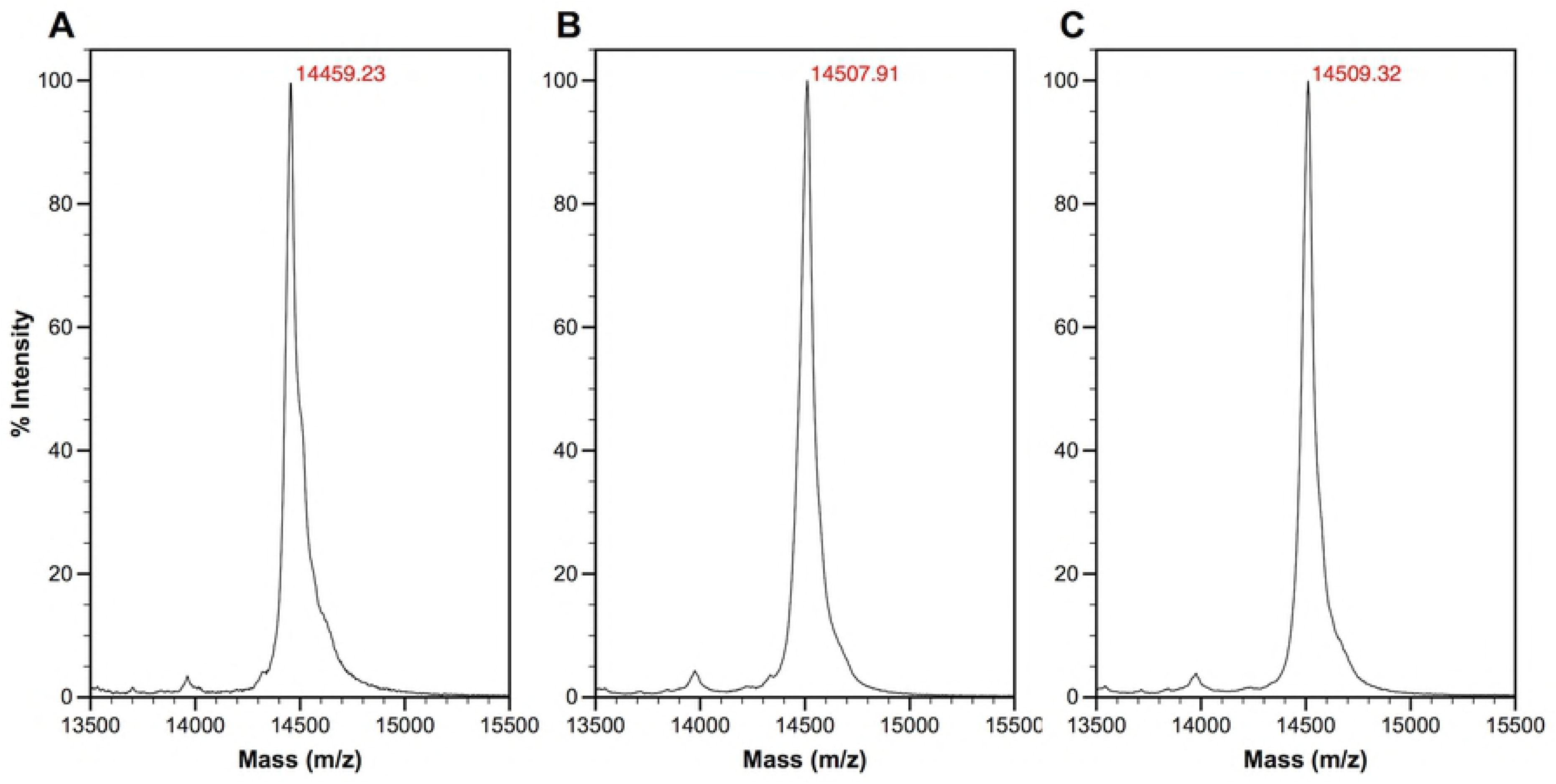
MALDI-TOF MS analysis of the N-terminal acetylation efficiency of pTSara-NatB. MALDI-TOF mass spectra of αSyn purified from *E. coli* transformed with pET21a-alpha-synuclein alone (A) or pET21a-alpha-synuclein+pTSara-NatB (B, C) and induced either only with 1 mM IPTG at OD_600_∼0.6 (A, C) or with 0.2% L-arabinose at OD_600_ ∼0.6, 30 min. before IPTG induction (B). The ∼42 Da shift in the intact mass of αSyn purified from co-transformed *E. coli* (predicted MW of N^α^-acetylated αSyn 14502.20 Da) shows how the basal constitutive expression of NatB (C) is sufficient to completely acetylate its overexpressed substrate.

Since MALDI-TOF MS could mask the presence of a small population of non-N^α^-acetylated substrate, trypsin digestion followed by MALDI-TOF MS was also performed on a control (non-N^α^-acetylated) sample and one of purified αSyn from non-L-arabinose-induced co-transformed *E. coli* (100% N^α^-acetylated according to MALDI-TOF MS). The mass spectrum of the fragments (Fig 3 and Table 1) confirms the N-terminal +42 Da mass shift that corresponds to N-terminal acetylation and the efficiency of the PTM (>97%).

**Table 1.**
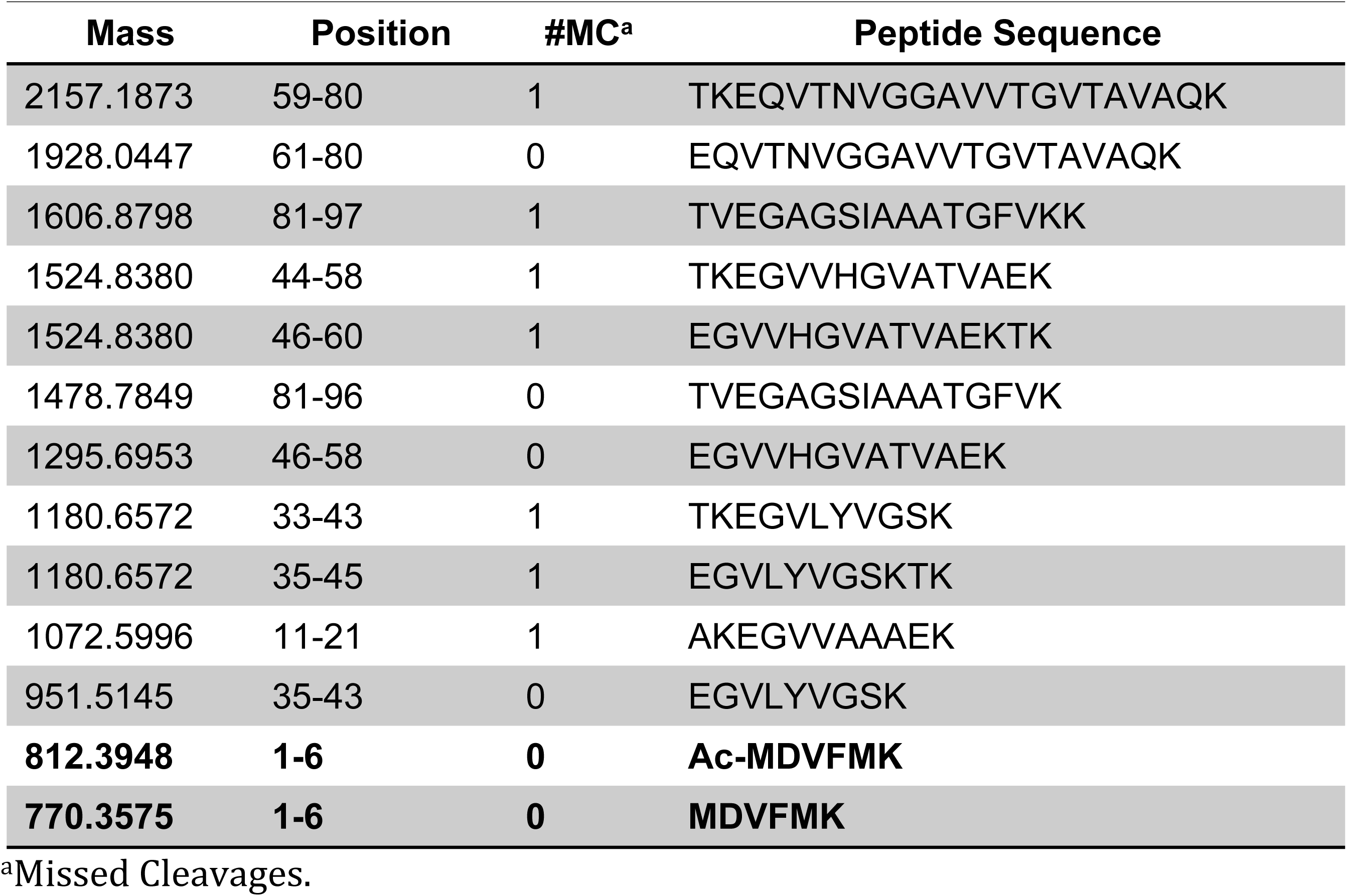
Identity of the most abundant peptide fragments identified in the MALDI-TOF mass spectra of trypsin-digested αSyn [27].

**Fig 3.**
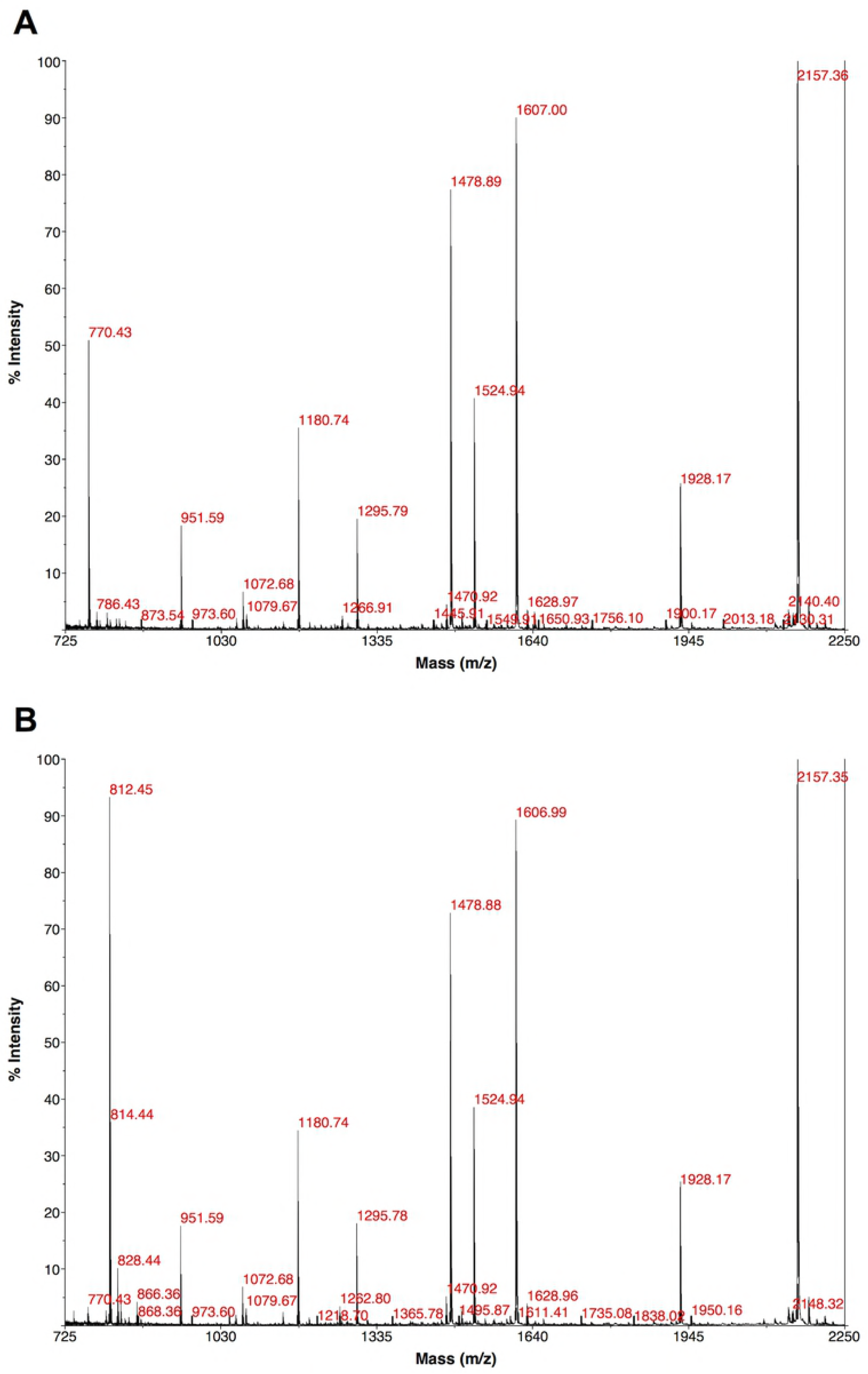
MALDI-TOF MS of trypsin-digested αSyn. MALDI-TOF mass spectra of trypsin-digested samples of αSyn purified from *E. coli* transformed with pET21a-alpha-synuclein alone (A) or pET21a-alpha-synuclein+pTSara-NatB (B) and only induced with 1 mM IPTG at OD_600_ ∼0.6. The ∼42 Da shift in the N-terminal fragment (770.43 Da → 812.45 Da, see Table 1) confirms the successful and complete N-terminal acetylation of αSyn upon NatB co-expression.

Finally, especially given the constitutive expression of our construct, the effects of widespread N-terminal acetylation on the bacterial proteome (and in particular its potential toxicity or metabolic modulation) were tested comparing the growth curves of BL21(DE3) transformed with either pET21a-alpha-synuclein alone or pET21a-alpha-synuclein+pTSara-NatB (with or without L-arabinose induction) (Fig 4). Both in non-L-arabinose-induced (pET+pTSara) and induced (pET+p TSara+0.2% ara) co-transformed *E. coli* there is an increase in the OD_600_ of the cultures, when compared to those of singly transformed bacteria (pET). No toxicity is thus observed, rather an increased bacterial proliferation (although the kinetics of the growth do not appear to be changed by the introduction of N-terminal acetylation), the reason of which has not been yet investigated. It must be noted how such increased culture density does not reflect in increased yields of N^α^-acetylated αSyn (in contrast with what previously reported for other NatB susbstrates [20]).

**Fig 4.**
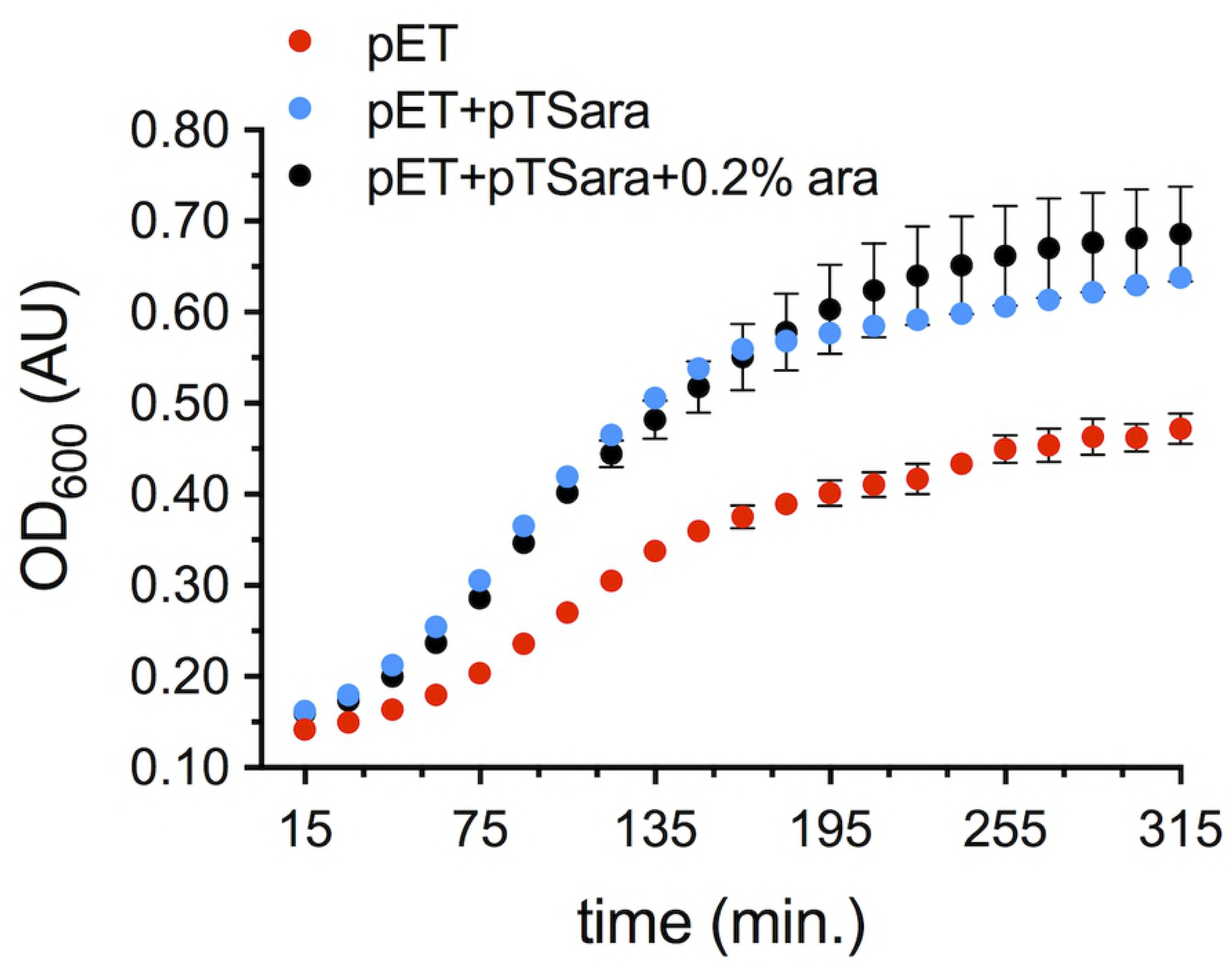
Impact of proteome-wide N-terminal acetylation on *E. coli* growth. Growth curves of *E. coli* transformed with pET21a-alpha-synuclein (pET) and grown in LB+amp or co-transformed with pET21a-alpha-synuclein+pTSara-NatB in the presence (pET+pTSara+0.2% ara) or absence (pET+pTSara) of 0.2% L-arabinose in LB+amp+cam. The absorbance (optical density) at 600 nm (OD_600_) is employed as a readout of their growth and the SDs obtained from 8 technical replicates are plotted.

## Discussion

In this work we developed and characterized pTSara-NatB, an improved N-terminal acetylation construct for recombinant protein expression in *E. coli*. We tested its ability to completely N^α^-acetylate our (model) target protein αSyn both upon L-arabinose induction and by relying only on the uninduced constitutive expression of the NatB complex subunits.

A clear advantage of the pTSara-NatB-mediated N-terminal acetylation is the ease which the uninduced, constitutive expression of NatB ensures. Viable substrates can be completely acetylated merely by the presence of the construct in co-transformed bacteria. In addition, the L-arabinose-inducible system (possibly in combination with a rhamnose-controlled expression vector in the case of problematic targets [21]) provides a flexibility that should secure the complete N-terminal acetylation of even intractable substrates. In addition, all our testing was done in LB medium and showed excellent N^α^-acetylation efficiency (pNatB has been reported to perform best in rich culture media, such as NZY [22]), which could possibly extend its use to minimal medium, as in the expression of isotopically-labeled recombinant proteins. We also observed, as previously reported [22], how N-terminal acetylation is complete only in freshly transformed *E. coli*.

Future developments should be the cloning of similar constructs for the other members of the NAT family [21], so to allow the recombinant expression of the whole N-terminal acetylome in bacteria, and extensive testing on a variety of substrates and culture conditions.

## Acknowledgments

We thank Zach Herbert, Jim Lee and the Molecular Biology Core Facilities of the Dana-Farber Cancer Institute for the use of instruments and assistance with Sanger DNA sequencing and mass spectrometry. We also thank our colleagues at the Ann Romney Center for Neurologic Diseases for many helpful discussions.

## Author Contributions

TB conceived and supervised the study; MR and TB designed experiments; MR, AEP, DSP and TB performed experiments; MR, AEP, DSP and TB analyzed data; MR and TB wrote the manuscript with input from AEP and DSP.

## Supporting Information

**S1 Fig. MALDI-TOF MS analysis of the N-terminal acetylation efficiency of pTSara-NatB in the presence of 0.2% D-glucose.** MALDI-TOF mass spectrum of αSyn purified from *E. coli* transformed with pET21a-alpha-synuclein+pTSara-NatB, grown in the presence of 0.2% D-glucose and induced with 1 mM IPTG at OD_600_ ∼0.6 (predicted MW of N^α^-acetylated αSyn 14502.20 Da).

**S2 Fig. Hydrophobic interaction chromatography (HIC) can resolve N^α^-acetylated and non-N^α^-acetylated αSyn mixtures.** (A) MALDI-TOF mass spectrum of αSyn purified from *E. coli* transformed with pET21a-alpha-synuclein+pNatB and induced with 1 mM IPTG at OD_600_ ∼0.6, showing a mixture of N^α^-acetylated and non-N^α^-acetylated αSyn. (B) Chromatogram of the HIC elution step of an aliquot from the same expression batch (in blue the 280-nm UV absorbance, in red the conductivity). HIC resolves N^α^-acetylated and non-N^α^-acetylated mixtures of αSyn (N^α^-acetylated αSyn has a slightly higher retention volume), as confirmed by MALDI-TOF MS on the two αSyn peaks, after size-exclusion chromatography (non-N^α^-acetylated αSyn, C; N^α^-acetylated αSyn, D).

**S3 Fig. Uncropped Western blots.**

## References

1. Aksnes H, Drazic A, Marie M, Arnesen T. First Things First: Vital Protein Marks by N-Terminal Acetyltransferases. Trends Biochem Sci. 2016;41: 746–760. doi:10.1016/j.tibs.2016.07.005

2. Liszczak G, Goldberg JM, Foyn H, Petersson EJ, Arnesen T, Marmorstein R. Molecular basis for N-terminal acetylation by the heterodimeric NatA complex. Nat Struct Mol Biol. 2013;20: 1098–105. doi:10.1038/nsmb.2636

3. Hong H, Cai Y, Zhang S, Ding H, Wang H, Han A. Molecular Basis of Substrate Specific Acetylation by N-Terminal Acetyltransferase NatB. Structure. Elsevier Ltd.; 2017;25: 641–649.e3. doi:10.1016/j.str.2017.03.003

4. Van Damme P, Evjenth R, Foyn H, Demeyer K, De Bock P-J, Lillehaug JR, et al. Proteome-derived Peptide Libraries Allow Detailed Analysis of the Substrate Specificities of N(alpha)-acetyltransferases and Point to hNaa10p as the Post-translational Actin N(alpha)-acetyltransferase. Mol Cell Proteomics. 2011;10: M110.004580. doi:10.1074/mcp.M110.004580

5. Gautschi M, Just S, Mun A, Ross S, Rücknagel P, Dubaquié Y, et al. The yeast N(alpha)-acetyltransferase NatA is quantitatively anchored to the ribosome and interacts with nascent polypeptides. Mol Cell Biol. 2003;23: 7403–14. doi:10.1128/MCB.23.20.7403-7414.2003

6. Polevoda B, Brown S, Cardillo TS, Rigby S, Sherman F. Yeast N(alpha)-terminal acetyltransferases are associated with ribosomes. J Cell Biochem. 2008;103: 492–508. doi:10.1002/jcb.21418

7. Bendor JT, Logan TP, Edwards RH. The function of α-synuclein. Neuron. Elsevier Inc.; 2013;79: 1044–66. doi:10.1016/j.neuron.2013.09.004

8. Iwai A, Masliah E, Yoshimoto M, Ge N, Flanagan L, de Silva HA, et al. The precursor protein of non-A beta component of Alzheimer’s disease amyloid is a presynaptic protein of the central nervous system. Neuron. 1995;14: 467–75. doi:10.1016/0896-6273(95)90302-X

9. Lautenschläger J, Kaminski CF, Kaminski Schierle GS. α-Synuclein - Regulator of Exocytosis, Endocytosis, or Both? Trends Cell Biol. 2017;27: 468–479. doi:10.1016/j.tcb.2017.02.002

10. Shulman JM, De Jager PL, Feany MB. Parkinson’s disease: genetics and pathogenesis. Annu Rev Pathol. 2011;6: 193–222. doi:10.1146/annurev-pathol-011110-130242

11. de Lau LML, Breteler MMB. Epidemiology of Parkinson’s disease. Lancet Neurol. 2006;5: 525–35. doi:10.1016/S1474-4422(06)70471-9

12. Anderson JP, Walker DE, Goldstein JM, de Laat R, Banducci K, Caccavello RJ, et al. Phosphorylation of Ser-129 is the dominant pathological modification of alpha-synuclein in familial and sporadic Lewy body disease. J Biol Chem. 2006;281: 29739–52. doi:10.1074/jbc.M600933200

13. Bartels T, Choi JG, Selkoe DJ. α-Synuclein occurs physiologically as a helically folded tetramer that resists aggregation. Nature. 2011;477: 107–10. doi:10.1038/nature10324

14. Trexler AJ, Rhoades E. N-Terminal acetylation is critical for forming α- helical oligomer of α-synuclein. Protein Sci. 2012;21: 601–5. doi:10.1002/pro.2056

15. Dikiy I, Eliezer D. N-terminal acetylation stabilizes N-terminal helicity in lipid- and micelle-bound α-synuclein and increases its affinity for physiological membranes. J Biol Chem. 2014;289: 3652–65. doi:10.1074/jbc.M113.512459

16. Bartels T, Kim NC, Luth ES, Selkoe DJ. N-alpha-acetylation of α-synuclein increases its helical folding propensity, GM1 binding specificity and resistance to aggregation. PLoS One. 2014;9: 1–10. doi:10.1371/journal.pone.0103727

17. Iyer A, Roeters SJ, Schilderink N, Hommersom B, Heeren RMA, Woutersen S, et al. The Impact of N-terminal Acetylation of α-Synuclein on Phospholipid Membrane Binding and Fibril Structure. J Biol Chem. 2016;291: 21110–21122. doi:10.1074/jbc.M116.726612

18. Selkoe D, Dettmer U, Luth E, Kim N, Newman A, Bartels T. Defining the native state of α-synuclein. Neurodegener Dis. 2014;13: 114–117. doi:10.1159/000355516

19. Jones JD, O’Connor CD. Protein acetylation in prokaryotes. Proteomics. 2011;11: 3012–22. doi:10.1002/pmic.201000812

20. Johnson M, Coulton AT, Geeves M a., Mulvihill DP. Targeted amino-terminal acetylation of recombinant proteins in E. coli. PLoS One. 2010;5: e15801. doi:10.1371/journal.pone.0015801

21. Eastwood T, Baker K, Brooker H, Frank S, Mulvihill DP. An enhanced recombinant amino-terminal acetylation system and novel in vivo high-throughput screen for molecules affecting α-synuclein oligomerisation. FEBS Lett. 2017;38: 42–49. doi:10.1002/1873-3468.12597

22. Johnson M, Geeves MA, Mulvihill DP. Production of amino-terminally acetylated recombinant proteins in E. coli. Methods Mol Biol. 2013;981: 193–200. doi:10.1007/978-1-62703-305-3-15

23. Guzman LM, Belin D, Carson MJ, Beckwith J. Tight regulation, modulation, and high-level expression by vectors containing the arabinose P(BAD) promoter. J Bacteriol. 1995;177: 4121–4130.

24. Shis DL, Bennett MR. Library of synthetic transcriptional AND gates built with split T7 RNA polymerase mutants. Proc Natl Acad Sci U S A. 2013;110: 5028–33. doi:10.1073/pnas.1220157110

25. Addgene: pNatB (pACYCduet-naa20-naa25) [Internet]. [cited 7 May 2018]. Available: https://www.addgene.org/53613/

26. Schleif R. AraC protein, regulation of the L-arabinose operon in Escherichia coli, and the light switch mechanism of AraC action. FEMS Microbiol Rev. 2010;34: 779–96. doi:10.1111/j.1574-6976.2010.00226.x

27. Wilkins MR, Lindskog I, Gasteiger E, Bairoch A, Sanchez JC, Hochstrasser DF, et al. Detailed peptide characterization using PEPTIDEMASS – a World-Wide-Web-accessible tool. Electrophoresis. 1997;18: 403–8. doi:10.1002/elps.1150180314

